# The Eya1 phosphatase mediates Shh-driven symmetric cell division of cerebellar granule cell precursors

**DOI:** 10.1101/668277

**Authors:** Daniel J. Merk, Pengcheng Zhou, Samuel M. Cohen, Maria F. Pazyra-Murphy, Grace H. Hwang, Kristina J. Rehm, Jose Alfaro, Xuesong Zhao, Eunyoung Park, Pin-Xian Xu, Jennifer A. Chan, Michael J. Eck, Kellie J. Nazemi, Rosalind A. Segal

**Author notes:** **Corresponding author**: Rosalind A. Segal, Dana-Farber Cancer Institute, 360 Longwood Avenue, Boston, MA 02215, USA.

## Abstract

During neural development, stem and precursor cells can divide either symmetrically or asymmetrically. The transition between symmetric and asymmetric cell divisions is a major determinant of precursor cell expansion and neural differentiation, but the underlying mechanisms that regulate this transition are not well understood. Here, we identify the Sonic hedgehog (Shh) pathway as a critical determinant regulating the mode of division of cerebellar granule cell precursors (GCPs). Using partial gain and loss of function mutations within the Shh pathway, we show that pathway activation determines spindle orientation of GCPs, and that mitotic spindle orientation directly correlates with the mode of division. Mechanistically, we show that the phosphatase Eya1 is essential for implementing Shh-dependent GCP spindle orientation. We identify atypical protein kinase C (aPKC) as a direct target of Eya1 activity and show that Eya1 dephosphorylates Threonine (T410) in the activation loop of this polarity complex component. Thus, Eya1 inactivates the cell polarity complex, resulting in reduced phosphorylation of Numb and other components that regulate the mode of division. This Eya1-dependent cascade is critical in linking spindle orientation, cell cycle exit and terminal differentiation. Together these findings demonstrate that a Shh-Eya1 regulatory axis selectively promotes symmetric cell divisions during cerebellar development by coordinating spindle orientation and cell fate determinants.

**Summary statement:** Biological responses to Shh signaling are specified by the magnitude of pathway activation and the cellular context. This study shows that potent Shh signaling regulates mitotic orientation and symmetric division of cerebellar granule cell precursors in a process that requires the phosphatase Eya1 and unequal distribution of cell fate determinants to daughter cells.

## Introduction

The morphogenic factor Hedgehog (Hh) was initially discovered in *Drosophila* in 1980 based on its role in segment polarity (Nüsslein-Volhard and Wieschaus, 1980). Hh has three vertebrate counterparts, and the closest homolog, Shh, functions as a key developmental regulator by acting as both a strong mitogen and a morphogen in multiple tissues (Ingham et al., 2011; Pak and Segal, 2016). Accordingly, it has long been appreciated that dysregulation of Shh signaling contributes to various forms of congenital disorders and cancer formation (McMahon et al., 2003; Nieuwenhuis and Hui, 2005). During cerebellar development, Shh acts as a critical mitogen for GCPs (Wechsler-Reya and Scott, 1999), and aberrant signaling of this pathway in GCPs results in Shh-subtype medulloblastoma (Taylor et al., 2012). In addition to acting as a strong mitogen for cerebellar and other neural progenitors, Shh signaling can also affect the mode of division in the developing cortex. Shh signaling has been suggested to promote symmetric divisions of cerebral cortical progenitors, and may also affect the mode of division in motor neuron precursors in the spinal cord (Araujo et al., 2014; Saade et al., 2017; Saade et al., 2013).

Changes in phosphorylation state are critical for most growth factor signaling pathways, including Shh signaling (Hillman et al., 2011; Nybakken et al., 2005; Purzner et al., 2018). Thus kinases including Protein kinase A (PKA), aPKC, as well as other phosphatases such as protein phosphatase 2A (PP2A) and Eya1 function as critical regulators of Shh signaling (Atwood et al., 2013; Eisner et al., 2015; Jia et al., 2009; Wang et al., 1999). The Eya1 phosphatase is a member of a small gene family originally described in *Drosophila*, where Eya has a key role in retina determination (Bonini et al., 1993). The mammalian homologues (*Eya1 − 4*) (Borsani et al., 1999; Xu et al., 1997; Zimmerman et al., 1997) play critical roles during development as highlighted by the fact that heterozygous loss of function mutations in *EYA1*, or its partner *SIX1*, are associated with the congenital disorder branchio-oto-renal syndrome, with ear, kidney and craniofacial defects (Abdelhak et al., 1997; Vincent et al., 1997). Eya1, Six1, and the closely related Eya and Six family members, are typically expressed at high levels during development and downregulated later in life. Not surprisingly, members of both families show aberrant re-activation in various tumor entities such as breast cancer (Wu et al., 2013), glioblastoma (Auvergne et al., 2013) and medulloblastoma (Cavalli et al., 2017).

All mammalian Eya proteins contain a highly conserved C-terminal Eya domain, which interacts with Six transcription factors or other members of the retinal determination network, and a less conserved N-terminal Eya domain (Tadjuidje and Hegde, 2013). Structurally, Eya1-4 belong to the superfamily of haloacid dehalogenases (HAD), which encompasses several different enzyme classes including dehalogenases, ATPases and magnesium-dependent phosphatases (Aravind et al., 1998). Surprisingly, Eya family members are reported to have dual protein Tyrosine (Tyr) as well as Serine(Ser)/Threonine(Thr) phosphatase activity (Li et al., 2017; Okabe et al., 2009; Sano and Nagata, 2011; Welcker et al., 2004). However, this view has recently been challenged by a study suggesting that Ser/Thr phosphatase activity of Eya3 might not be intrinsic, but rather arises from association with the PP2A (Zhang et al., 2018). Moreover, since Eya acts as both a co-transcriptional activator and a phosphatase, it is not clear whether the intrinsic phosphatase activity is required for its actions (Ahmed et al., 2012; Li et al., 2003; Wu et al., 2013). In previous studies we provided evidence that Eya1 phosphatase activity is essential to promote Shh signaling (Eisner et al., 2015). While several other studies also suggest that phosphatase activity contributes to Eya-dependent transcriptional output (Davis et al., 2017), data from the *Drosophila* system suggest that phosphatase activity of the Eya family members may be dispensable for many of its actions (Jemc and Rebay, 2007; Jin et al., 2013; Tootle et al., 2003). Thus far, four physiological substrates of Eya phosphatase activity have been identified: the histone H2A variant H2AX (Cook et al., 2009; Krishnan et al., 2009), the estrogen receptor ERβ (Yuan et al., 2014), the proto-oncogene Myc (Xu et al., 2014) and Notch1 (Zhang et al., 2017), and additional potential substrates have been suggested (El-Hashash et al., 2011; El-Hashash et al., 2017; Li et al., 2017). It is not clear, which, if any, of these substrates might be important for Shh signaling.

In this study, we address novel roles of Shh signaling and Eya1 activity in regulating GCP development. We show that Shh signaling regulates spindle orientation of mitotic GCPs during cerebellar development and that spindle orientation directly correlates with the mode of division. We further show that Eya1 is required for the effects of Shh signaling on GCP spindle orientation. We find that aPKC is a direct substrate of Eya1, and that de-phosphorylation of aPKC ζ by Eya1 leads to hypo-phosphorylation of the cell fate determinant Numb, a well-known target of aPKC ζ In this way, Eya1 enables Shh signaling to drive symmetric cell division of cerebellar GCPs during early postnatal development. Together these data provide mechanistic insight into Shh pathway regulation of symmetric cell division and the rapid expansion of the developing GCP population.

## Results

### Shh regulates spindle orientation of proliferating GCPs

Previous studies suggest that the plane of cell division for GCPs is developmentally regulated (Haldipur et al., 2015; Zagon and McLaughlin, 1987), with a progressive increase of mitotic figures with the plane of division oriented horizontally to the pial surface during postnatal development. However, a more recent study instead suggested that vertical divisions are predominant at late postnatal stages of GCP development (Miyashita et al., 2017). Therefore, we first determined the plane of dividing GCPs during the peak of proliferation (postnatal day 1 (P1) to P10) in the murine cerebellum by visualizing anaphase cells using antibodies to phospho-Histone H3. During postnatal development, there is a gradual increase of the proportion of GCP cell divisions with a horizontal mitotic plane (i.e. parallel to the pial surface) (Fig. S1A, B). Further analyses showed that mitotic orientation did not correlate with the distance from the pial surface, ruling out the possibility that the plane of cleavage might merely be regulated by the actual depth of the dividing cell within the outer external granule cell layer (Fig. S1C). These data demonstrate that GCPs transition from vertical to horizontal divisions during the period of early postnatal development when Shh regulates proliferation (Wechsler-Reya and Scott, 1999).

As recent data suggest that Shh signaling can directly promote symmetric divisions of neuroepithelial cells (Saade et al., 2017), we carefully assessed spindle orientation of GCPs in mouse models with reduced or enhanced Shh signaling. To do so, we used antibodies for γ-tubulin and acetylated α-tubulin to simultaneously visualize centrioles and spindles, respectively, within thick sections of early cerebellar cortex (Fig. 1A, B). Mice expressing a mutant form of Shh (*Shh^Ala^*) with impaired proteoglycan interactions exhibit decreased GCP proliferation due to reduced Shh output (Chan et al., 2009). Here we show that this hypomorphic mutation alters mitotic spindle orientation, with a decrease in vertical and an increase in horizontal cell divisions of GCPs at P3 (Fig. 1C). In striking contrast to this partial loss of function phenotype, we find that Shh pathway gain of function *Ptch1*^+/−^ mice (Goodrich et al., 1997) show a significant increase in vertical cell divisions in the external granule cell layer (EGL) (Fig. 1D). Together, these gain and loss of function studies demonstrate that the Shh pathway both regulates spindle orientation and promotes cell cycle progression of cerebellar GCPs.

**Fig. 1.**
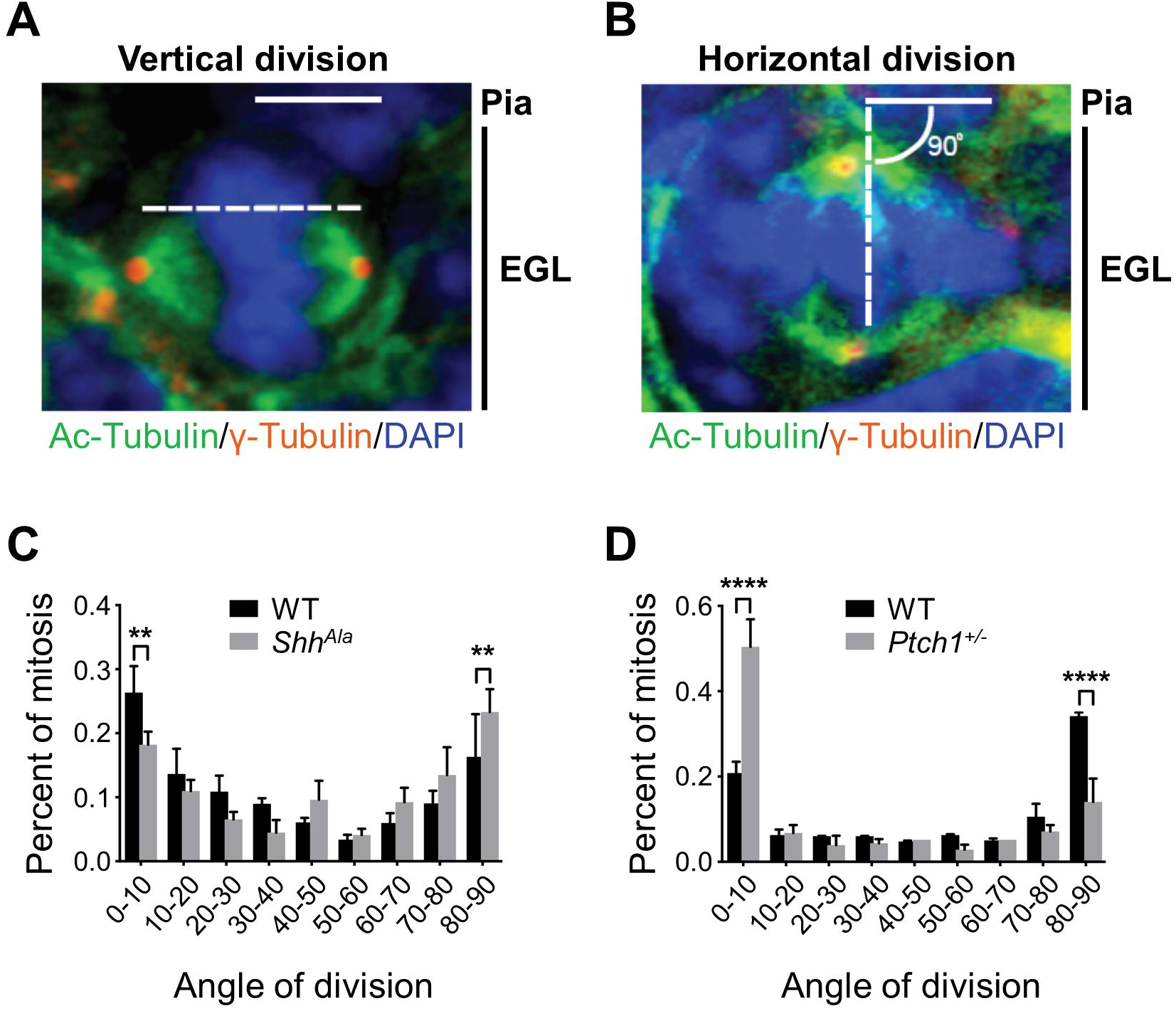
Shh signaling regulates spindle orientation of mitotic GCPs. Representative immunostainings at P3 visualizing both centrioles (γ-Tubulin) and spindles (Ac-Tubulin) of mitotic GCPs in anaphase with the plane of division oriented vertically (A) and horizontally (B) with respect to the pial surface. Straight line represents the pial surface, broken line represents the plane of division as determined by the position of the centrioles. (C, D) Bar graphs illustrating the distribution of angles of division of mitotic GCPs in the EGL of *Shh^Ala^* mice (C) or *Ptch1*^+/−^ mice (D) as compared to wild type (WT) mice (n = 5 for each condition). \*\**P*<0.01, \*\*\*\**P*<0.0001, Fisher’s exact test. Data are mean±s.e.m.

### Spindle orientation correlates with GCP cell fate

While GCPs uniformly give rise to mature granule cell neurons, the progenitors undergo migration and terminal differentiation over a prolonged time period. Thus, in the early postnatal period, some GCPs migrate from the EGL to the inner granule cell layer (IGL) and differentiate, while other GCPs re-enter the cell cycle, providing two distinct fates for sister cells after each mitotic division. To determine whether mitotic orientation correlates with symmetrical or asymmetrical cell fates of sister cells, we carried out real time analysis of sparsely labeled GCPs in organotypic slice cultures. To do so, we generated *Atoh1-creER::GFP*^*Fl*/+^ mice and injected those mice with a low dose of tamoxifen to selectively label a subset of developing GCPs (Machold and Fishell, 2005), then prepared organotypic brain slices and monitored individual dividing cells and their corresponding daughter cells over time (Fig. 2). We followed 57 GFP-positive, dividing GCPs from P3 and 31 from P6. Consistent with our previous data, we observed more vertical divisions at P3 (P3: 34 vertical, 12 horizontal, 11 oblique) as compared to P6 (P6: 10 vertical, 18 horizontal, 3 oblique) (p=0.0022, χ^2^ test). Further analyses of these images suggest that the fates of daughter cells differ depending on the spindle orientation. After vertical division, both daughter cells remain in the EGL (34/34 at P3, 10/10 at P6). In contrast, in many horizontal divisions (6/12 at P3 and 5/18 at P6), one of the daughter cells remains in the EGL while the other one migrates into the IGL (p=0.001). These functional studies support the hypothesis that vertically aligned spindles correlate with symmetric cell division, whereas horizontal divisions generate two daughter cells with distinct cell fates.

**Fig. 2.**
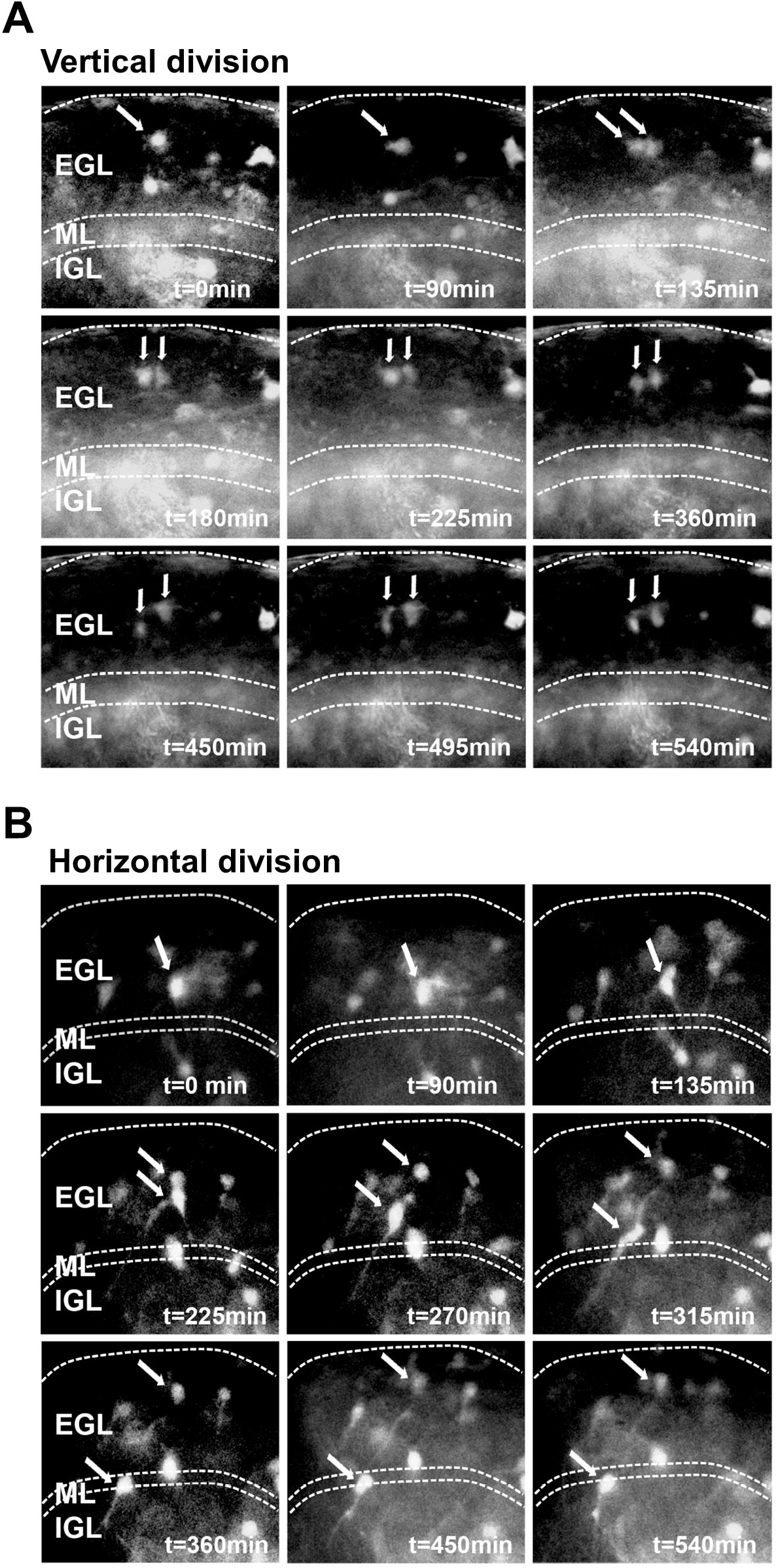
Spindle orientation of dividing GCPs correlates with migrational direction of daughter cells. (A, B) Representative images of individual, GFP-labeled GCPs followed over a period of 9 hours. Single arrows indicate cells before mitosis and double arrows indicate position of daughter cells. Dotted lines indicate boundaries of the different layers of the cerebellar cortex. In vertical divisions (A), both daughter cells remain in the EGL. In contrast, in horizontal divisions (B), one daughter cell remains in the EGL while the other one migrates into the IGL.

While our data argue for a mechanism that links spindle orientation and mode of division of GCPs, our study does not demonstrate whether daughter cells undergoing horizontal division are distinct from one another at the time of mitosis, or acquire different fates after cell division is complete, perhaps due to changes in the microenvironment (Choi et al., 2005; Leto et al., 2016). The Par complex, consisting of aPKC, Par-3 and Par-6, has a highly conserved function in regulating spindle orientation and symmetric cell division (Durgan et al., 2011; Hao et al., 2010), and cells that are asymmetric at the time of division frequently show unequal distribution of multiple intracellular components that interact directly with the Par complex (Peyre et al., 2011; Rhyu et al., 1994). We therefore tested the possibility that Shh regulation of spindle orientation and cell fate in GCPs might involve unequal distribution of two Par complex partners, the cell fate determinant Numb and the spindle regulator LGN (Fig. 3A). We developed a quantitative symmetry index (Fig. 3B) to analyze the distribution of Numb and LGN in dividing, anaphase GCPs in which the mitotic orientation was either vertically or horizontally aligned to the pial surface. While Numb (Fig. 3C, D) and LGN (Fig. 3E, F) are distributed equally in cells with the plane of division vertical to the pial surface, these markers are instead distributed unequally in cells with the plane of division horizontal to the pial surface. Of note, we did not find any difference in the distribution of a control protein, GAPDH, with respect to mitotic orientation (Fig 3G, H). Taken together, these data indicate that mitotic spindle orientation directly correlates with the mode of division of proliferating GCPs, through a mechanism that involves the Par complex and regulated intracellular distribution of key interacting partners.

**Fig. 3.**
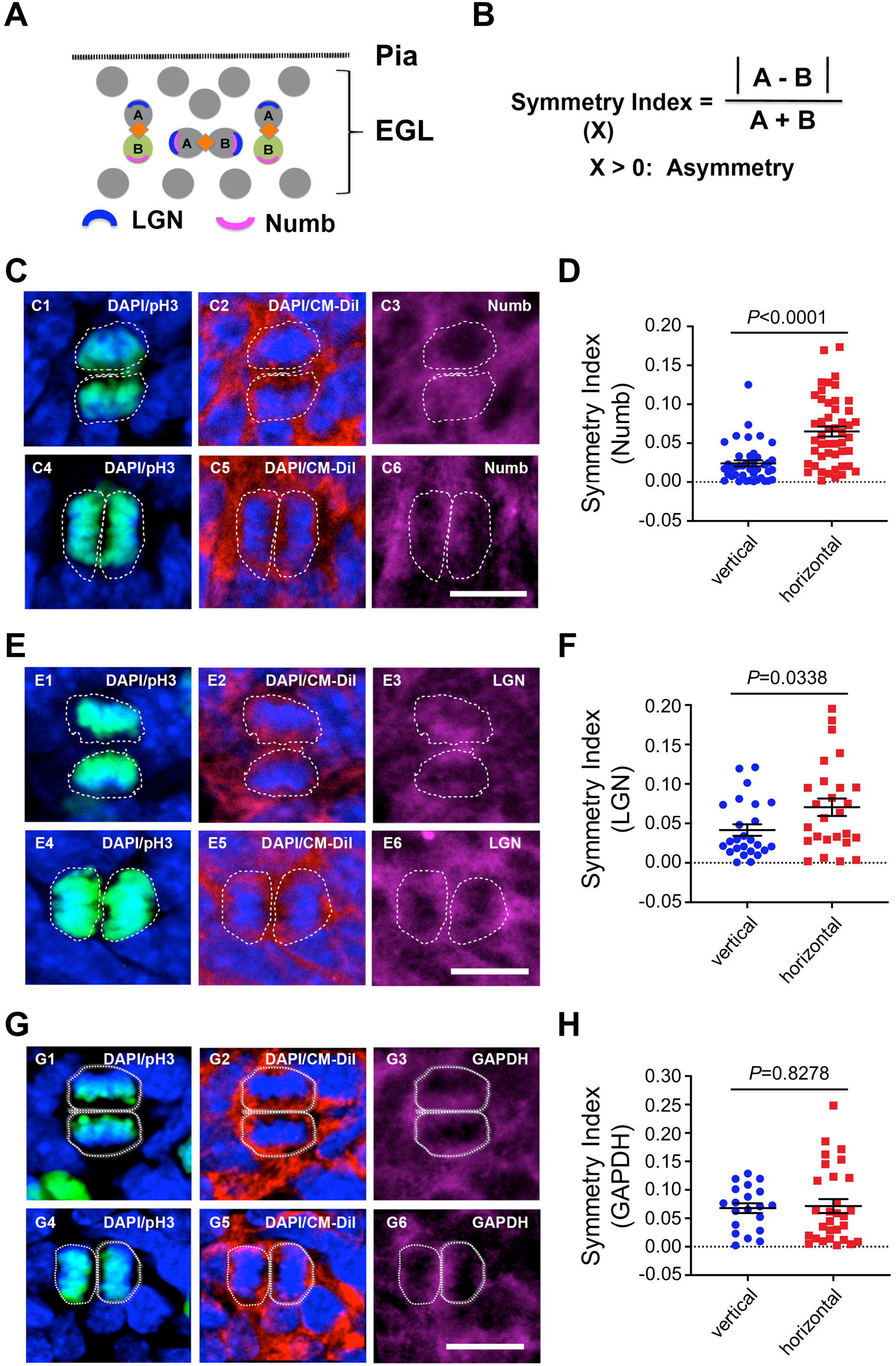
Spindle orientation of dividing GCPs correlates with the distribution of symmetry markers. (A) Schematic of GCP division pattern and protein distribution within daughter cells. (B) Formula for generating a symmetry index. (C-H) Representative immunostainings in cerebellar sections from wild type mice at P3 for Numb (C), LGN (E), and GAPDH (G) and corresponding quantitative analyses (D, F, H) for the degree of symmetric distribution. Distribution of cellular markers was quantified in GCPs dividing horizontally (C1-C3, E1-E3, G1-G3) or vertically (C4-C6, E4-E6, G4-G6) to the pial surface. Cellular markers were co-visualized with phospho-Histone H3 (pH3), CM-Dil and DAPI. White dashed lines represent the cell boundaries visualized using CM-Dil. Significance was determined using the unpaired *t* test. Scale bars, 10 μm.

### Eya1 is essential for Shh-dependent promotion of symmetric GCP divisions

We have previously described the phosphatase Eya1 as a positive regulator of Shh signaling during cerebellar development (Eisner et al., 2015). Consistent with this role, we find that heterozygous loss of *Eya1* in GCPs (Xu et al., 1999) reduced expression of known Shh pathway members and downstream targets, whereas expression of the Shh ligand itself was not affected (Fig. 4A). Strikingly, expression of *Eya1* closely correlates with expression of the major Shh target *Ptch1* over the course of cerebellar development (Fig. 4B), with peak levels of expression at postnatal day 6. Together these data indicate that Eya1 regulates the Shh transduction process rather than ligand expression. To determine whether Eya1 enables Shh-dependent orientation of mitotic GCPs, we examined spindle orientation in *Eya1*^+/−^ mice at P6. We visualized proliferating GCPs in the EGL using antibodies against phospho-Histone H3 and assessed the angle of division relative to the pial surface for anaphase cells (Fig. 4C). The dividing GCPs in the heterozygotes exhibited a significantly lower proportion of vertical divisions and higher proportion of horizontal divisions as compared to wild type controls (*P* = 0.045, χ^2^ test) (Fig. 4D). Thus *Eya1*^+/−^ mice phenocopy the changes in GCP spindle orientation observed in the hypomorphic, *Shh^Ala^* mice (Fig. 1C). Consistent with our previous data examining the spindle directly, *the Ptch1*^+/−^ gain of function mice exhibit increased vertical GCP cell divisions compared to wild type animals due to enhanced Shh pathway activation (*P* = 0.002). To identify genetic interactions between the Eya1 and Shh pathway mutations, we generated mice that are both heterozygous for *Eya1* and *Ptch1*. Loss of one copy of *Eya1* in the *Eya1*^+/−^ *Ptch1*^+/−^ mice abrogated the increase of vertical divisions observed in *Ptch1*^+/−^ mice (*P* = 0.9653), and restored the wild type proportions of vertical and horizontal divisions. This genetic interaction suggests that Eya1 is essential for the ability of Shh signaling to promote vertical, symmetric cell divisions of GCPs.

**Fig. 4.**
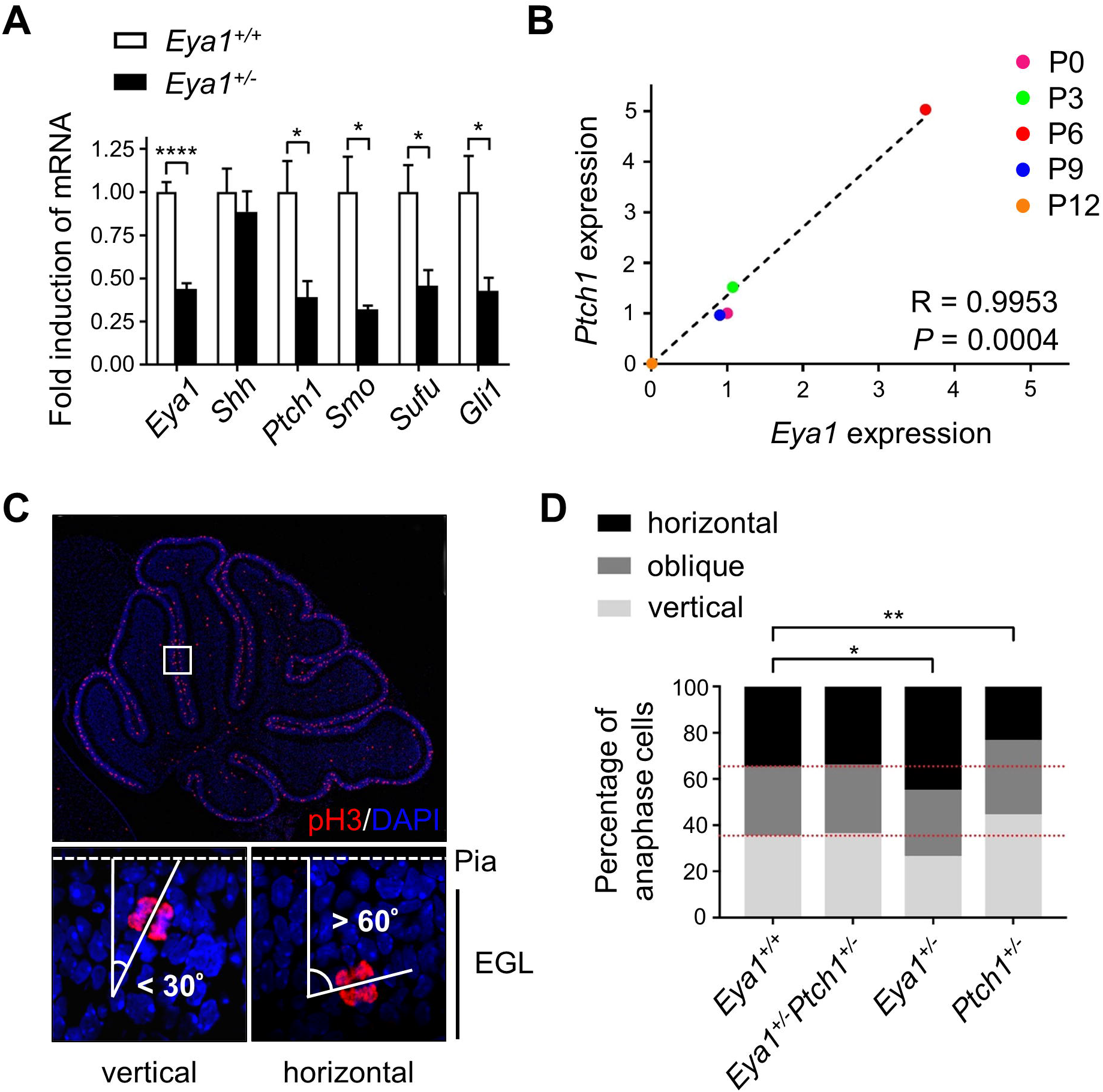
Expression of Eya1 enables Shh-dependent regulation of spindle orientation. (A) Quantitative real time PCR analyses for *Eya1* and Shh pathway members in P6 cerebellar lysates wild type controls and mice carrying a heterozygous knockout of *Eya1*. \**P*<0.05, \*\*\*\**P*<0.0001, unpaired *t* test. Data are mean±s.e.m. (B) Dot plot illustrating the correlation of expression of *Eya1* and *Ptch1* over the course of early development. Pearson correlation was computed. (C) Immunostainings for phospho-Histone H3 (pH3) of cerebellar sections from P6 wild type mice and representative images for vertically and horizontally dividing GCPs in the EGL. (D) Distribution of angles of division of mitotic GCPs in the EGL of *Eya1*^+/+^, *Eya1*^+/−^ *Ptch1*^+/−^, *Eya1*^+/−^ and *Ptch1*^+/−^ mice (n = 5 for each genotype). Angles were defined as being vertical (0-30°), oblique (30-60°), or horizontal (60-90°). \**P*<0.05, \*\**P*<0.01, χ^2^ test.

### Eya1 dephosphorylates atypical protein kinase C and regulates Numb phosphorylation

It is possible that the loss of function phenotype in *Eya1*^+/−^ mice is merely due to an overall impairment of Shh signaling, or Eya1 may function directly to regulate mode of division of GCPs. We have shown here that the intracellular distribution of the cell fate determinant Numb (Le Borgne et al., 2005; Schweisguth, 2004) correlates with GCP spindle orientation. Unequal distribution of Numb during mitosis is known to be a major regulator of asymmetric division, and phosphorylation by the Par complex component aPKC dictates intracellular localization of Numb during asymmetric division and migration (Smith et al., 2007; Zhou et al., 2011). We therefore investigated whether Eya1 might be involved in directly regulating the mode of division of GCPs by altering the aPKC/Numb axis, consistent with previous suggestions (El-Hashash et al., 2011), a study that has subsequently been retracted (El-Hashash et al., 2017). As mice with a homozygous knockout of *Eya1* are not viable at postnatal stages (Xu et al., 1999), we first used mouse embryonic fibroblasts (MEFs) from *Eya1*^−/−^ mice to study the effects of a complete loss of this phosphatase on both aPKC and Numb phosphorylation. Of note, Shh pathway activation in wild type MEFs using the Smoothened agonist SAG (300nM) confirmed induction of Gli transcription factors (Fig. S2), indicating that MEFs are susceptible to Shh activation. We carried out western blot analyses using an antibody against PKC with phosphorylation of the activation loop site; this antibody recognizes a conserved site that is constitutively phosphorylated by PDK1 in all PKC enzymes. We find that there is an increase in phosphorylation state at this site in *Eya1*^−/−^ cells compared with wild type MEFs (Fig. 5A, B). A more selective antibody that recognizes phosphorylation at the activation loop in one of the atypical PKCs, aPKC ζ (pT410), also showed enhanced phosphorylation in *Eya1*^−/−^ MEFS. In contrast, Eya1 mutations did not alter the phosphorylation state at T560, the turn motif site in aPKC ζ (Chou et al., 1998), suggesting that Eya1 may have a specific effect on the activation loop phosphorylation state of aPKC. While phosphorylation in the activation loop can affect both protein stability and enzymatic activity, we did not detect any changes in total levels of aPKC ζ in the *Eya1*^−/−^ MEFs. Taken together these findings indicate that the phosphatase Eya1 directly, or indirectly, regulates the phosphorylation state of the activation loop in PKC proteins, particularly in atypical PKCs, but these changes do not alter total levels of PKC.

**Fig. 5.**
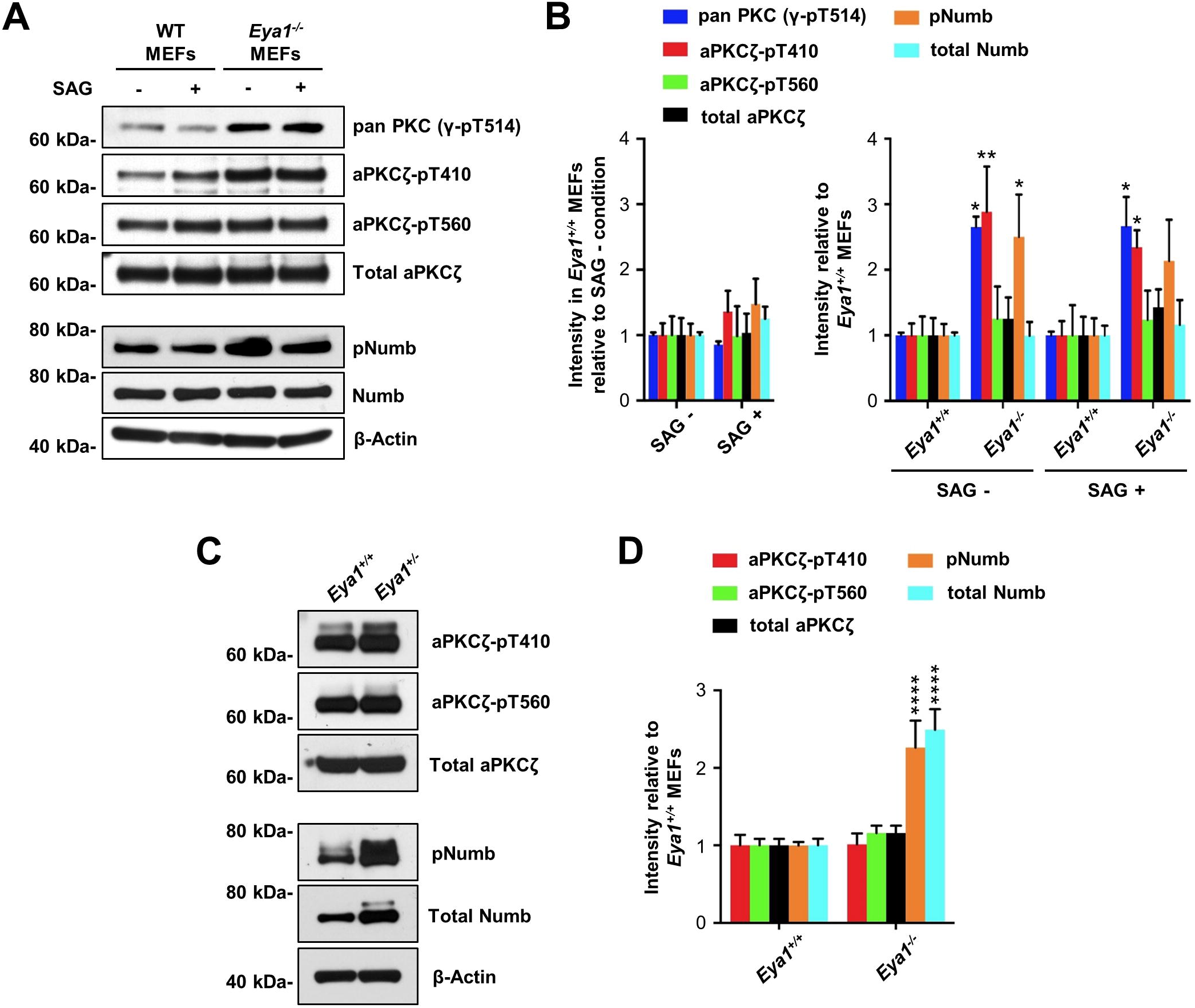
Eya1 regulates aPKCζ and Numb phosphorylation. (A) Representative western blots of protein lysates from MEF cells generated from wild type or *Eya1*^−/−^ mice, using indicated phospho-specific antibodies for protein kinase C (PKC) and Numb, and corresponding quantification (B) of relative protein levels from independent experiments (n = 4), Actin was used as a loading control. \**P*<0.05, \*\**P*<0.01, Two-way ANOVA with Bonferroni *post hoc* test. Data are mean±s.e.m. (C) Representative western blots of P6 cerebellar protein lysates from *Eya1*^+/+^ and *Eya1*^+/−^ mice using phospho-specific antibodies for aPKCζ and Numb, and corresponding quantification (D) of relative protein levels from independent experiments (n = 4), Actin was used as a loading control. \*\*\*\**P*<0.01, Two-way ANOVA with Bonferroni *post hoc* test. Data are mean±s.e.m.

Numb protein is a critical substrate of aPKC that has been implicated in spindle orientation and asymmetric cell division. We found that phosphorylation of Numb at the aPKC site was also significantly increased in the absence of *Eya1*. Again, these changes in phosphorylation state were not associated with any changes in the overall levels of Numb. Of note, Shh-pathway activation by SAG treatment alone had no significant effect on aPKC and Numb phosphorylation, suggesting that these changes were due to *Eya1* function and not due to general changes in Shh signaling.

As Eya1 affects both mitotic orientation and Shh signaling in the developing cerebellum, we analyzed aPKC ζ and Numb phosphorylation in cerebellar lysates from wild type and *Eya1*^+/−^ mice. Although we did not observe any changes in aPKC phosphorylation in these heterozygotes, we did observe increased phosphorylation of Numb after heterozygous loss of *Eya1* (Fig. 5C, D). In addition, we observed an increase in total Numb protein levels. These data suggest that the phosphatase Eya1 regulates the phosphorylation state of the cell fate determinant Numb, and that this change in phosphorylation may also affect protein stability.

Changes in the phosphorylation state of Numb in Eya1 mutants could reflect direct or indirect effects of the Eya1 phosphatase. As Eya1 binds to its substrates to initiate phosphatase activity, we used HA antibodies to immunoprecipitate protein lysates from HEK293T cells expressing HA-tagged Eya1, and then blotted for aPKC ζ and Numb (Fig. 6A). Immunoblotting for Six1, a well-known interaction partner of Eya1 phosphatase (Buller et al., 2001), was included as a positive control. Both Six1 and aPKC ζ protein were co-immunoprecipitated with HA-Eya1. In contrast, Numb protein was not present in the co-immunoprecipitates. We next performed *in vitro* phosphatase assays with purified, full length, recombinant wild type Eya1 protein that was expressed and purified from S2 cells, and various phospho peptides corresponding to sequences from aPKC ζ, Numb and H2AX as substrates (Fig. 6B). Purified wild type Eya1 catalyzed the dephosphorylation of a peptide corresponding to the known phospho Tyr motif from the Eya1 substrate H2AX (Cook et al., 2009). The D327 residue in the Eya domain has previously been shown to be required for enzymatic activity, and a D327A substitution results in a phosphatase-dead mutant protein (Cook et al., 2009; Li et al., 2017; Tootle et al., 2003). While wild type Eya1 showed a dose-dependent activity profile in the *in vitro* assay, Eya1 D327A protein showed essentially no activity even at high molar concentrations (Fig. 6C). Using this *in vitro* phosphorylation assay, we tested phospho Thr and phospho Ser peptides corresponding to the phosphorylation sites in aPKC and Numb that are altered in *Eya1* mutants *in vitro* and/or *in vivo*. While Eya1 showed marked activity using the phospho Thr aPKC ζ-T410 peptide (Fig. 6D), corresponding to the activation loop of aPKC ζ (Parker and Parkinson, 2001), purified Eya1 did not dephosphorylate the phospho Thr aPKC ζ-T560 peptide from the turn motif of the enzyme. Strikingly, Eya1 exhibited little or no activity on peptides corresponding to phospho Ser sites pS7, pS276, and pS295 of Numb (Smith et al., 2007). While wild type Eya1 displayed a dose-dependent activity profile for the aPKC ζ-T410 peptide, Eya1 D327A protein showed a significant reduction in phosphatase activity (Fig. 6E), consistent with previous studies showing that the C-terminal domain of Eya1 is essential for Thr and Tyr phosphatase activity (Li et al., 2017). These studies suggest that aPKC (T410) is a direct substrate of Eya1, but that Numb is not, and that changes in Numb instead reflect altered phosphorylation by aPKC.

**Fig. 6.**
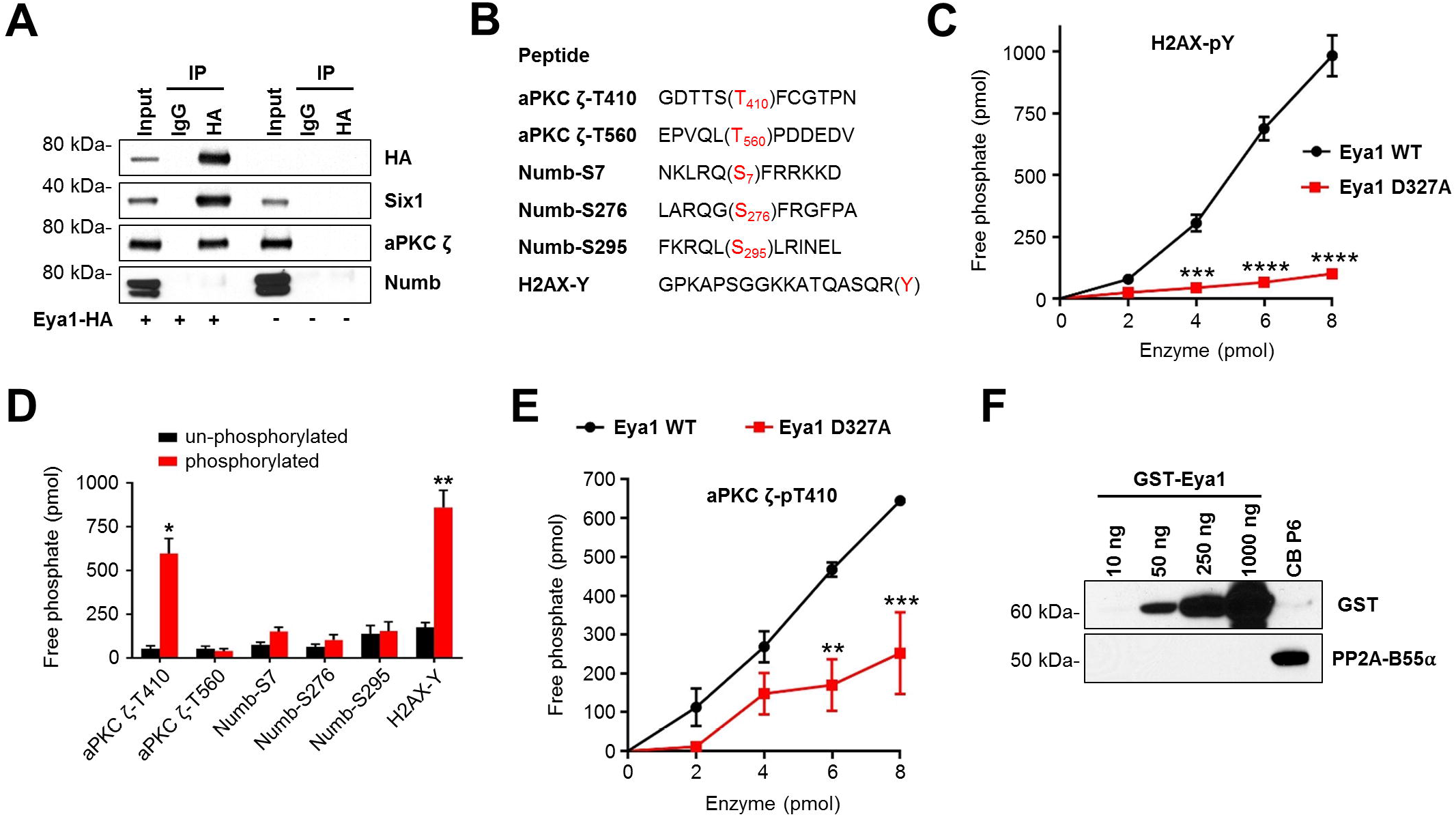
aPKC ζ is a direct substrate of intrinsic Eya1 threonine phosphatase activity. (A) HEK293T cells were transfected with Eya1-HA and protein lysates were precipitated with anti-HA antibody. Immunoprecipitates were blotted with anti-HA, anti-Six1, anti-aPKC ζ, and anti-Numb antibodies. Un-transfected HEK293T cells were included as a negative control. (B) Sequences of phospho-peptides used for *in vitro* Eya1 phosphatase assays. Phosphorylated threonine (T), serine (S) and tyrosine (Y) residues are highlighted in red. (C) Dose-response of phosphatase activity generated by wild type Eya1 enzyme or a mutated D327A form using H2AX-pY phosphorylated peptides as substrate (n = 3). \*\*\**P*<0.001, \*\*\*\**P*<0.0001, Twoway ANOVA with Bonferroni *post hoc* test. Data are mean±s.e.m. (D) Bar graphs representing the amount of free phosphate generated by Eya1 phosphatase activity using the indicated un-phosphorylated or phosphorylated peptides (n = 4). \**P*<0.05, \*\**P*<0.01, paired *t* test. Data are mean±s.e.m. (E) Dose-response of phosphatase activity generated by wild type Eya1 enzyme or a mutated D327A form using aPKC ζ-pT410 phosphorylated peptides as substrate (n = 3). \*\**P*<0.01, \*\*\**P*<0.001, Twoway ANOVA with Bonferroni *post hoc* test. Data are mean±s.e.m. (F) Western blot analysis of GST-tagged Eya1 protein isolated from S2 insect cells. Purified protein was blotted with anti-GST and anti-PP2A-B55α antibodies. Protein lysate (20 μg) from P6 cerebellum was included as a positive control for PP2A-B55α protein.

It had been suggested that the Ser/Thr phosphatase activity of Eya3 is not intrinsic but stems from its association with the protein phosphatase 2A (PP2A)-B55α holoenzyme (Zhang et al., 2018). As we hypothesize that phosphorylation state of aPKC depends on Eya1 activity, we asked whether PP2A-B55α is essential for this function. To do so, we checked for the presence of PP2A-B55α in the recombinant GST-tagged Eya1 prepared from S2 cells. While the Eya1 preparation showed a clear dose-dependent increase in reactivity with the GST-specific antibody, we did not detect any signal for PP2A-B55α protein at any concentration of Eya1 protein tested (Fig. 6F). In contrast, we readily detected PP2A-B55α protein in P6 cerebellar lysates. As PP2A-B55α protein is not present in detectable amounts in our enzymatically active Eya1 protein preparations, the phosphatase activity that dephosphorylates aPKC ζ appears to be an intrinsic feature of Eya1 that is altered in D327A mutant. Together, these data indicate that Eya1 directly and specifically dephosphorylates aPKC at residue T410, thereby reducing aPKC-mediated phosphorylation of Numb, altering Numb distribution and regulating spindle orientation and asymmetric cell division in developing GCPs.

## Discussion

The results presented here indicate that the Shh pathway and its positive effector Eya1 regulate spindle orientation and the mode of division of GCPs by promoting vertical spindle orientation and symmetric cell divisions. This finding is consistent with previous data showing that active Shh signaling promotes symmetric, self-renewing cell divisions of neuronal progenitors in multiple brain regions (Araujo et al., 2014; Haldipur et al., 2015; Saade et al., 2017; Saade et al., 2013; Yang et al., 2015). We provide both mechanistic and functional insight into Shh-regulated spindle orientation and symmetric division.

Previous studies suggest that Shh signaling drives symmetric neuroepithelial cell division by a process involving recruitment of protein kinase A to the centrosomes that nucleate the mitotic spindle (Saade et al., 2017). Our study adds to these findings by showing that the Eya1 phosphatase promotes Shh pathway-dependent mitotic spindle orientation of GCPs. Interestingly, we found that changes of both Shh signaling and/or *Eya1* expression seem to elicit directed turns from vertical to horizontal divisions or vice versa, rather than leading to progressive shift of the angles of division of mitotic GCPs. This indicates that proliferating GCPs actively decide between two regulated states of spindle orientation in a rather binary fashion. Furthermore, our data shows that spindle orientation of mitotic GCPs correlates with the mode of division. Real-time monitoring of proliferating GCPs revealed that spindle orientation is a strong indicator whether both daughter cells remain in the EGL (symmetric division) or one of the daughter cells will migrate into the IGL while the other one remains in the EGL (asymmetric division). Finally, unequal distribution of the cell-fate determinant Numb and the vertebrate homologue of Pins (LGN) further supports this model and provides evidence that GCPs begin to adopt a particular fate during mitosis. Similarly, in other epithelia such as the skin, mitotic orientation and mode of division are tightly linked, and precursors use symmetric and asymmetric division to generate appropriate ratios between proliferation and differentiation (Lechler and Fuchs, 2005; Williams et al., 2011).

Previous data have implicated Eya in Hh signaling in *Drosophila* and have provided multiple possible models for the actions of Eya family members in Shh signaling in vertebrates (Eisner et al., 2015; Lu et al., 2013; Pappu et al., 2003). Here, we show that Eya1 phosphatase activity can remove phosphates from the Thr site within the activation loop of aPKC ζ. After genetic loss of *Eya1* or developmental downregulation of *Eya1* expression, activated aPKC phosphorylates the endocytic component Numb, a change that can facilitate asymmetric cell division (Smith et al., 2007). Accordingly, during early development of GCPs, high expression of *Eya1* and active Shh signaling drive symmetric cell divisions. The data presented here suggest that the Eya1/aPKC/Numb axis links Shh pathway output to the cellular polarity machinery and facilitates regulation of the mode of division in response to a mitogenic cue. Previous studies indicate that active aPKC can regulate spindle orientation and polarize the intracellular distribution of Numb. In mammalian epithelia, Par proteins bind to aPKC ζ and LGN to form an apically located complex which establishes cell polarity and regulates spindle orientation (Durgan et al., 2011; Hao et al., 2010). aPKC itself is essential for regulation of spindle orientation in a process conserved across species (Guilgur et al., 2012). Data that aPKC is a direct substrate for Eya1 phosphatase activity indicate that Eya1 contributes both to regulation of cell polarity/spindle orientation and to cell fate determination. Interestingly, we have previously shown that aPKC activation and downstream Numb phosphorylation is an important mechanism to promote chemotaxis and migration of GCPs (Zhou et al., 2011), and differentiation of GCPs is tightly linked to their migration out of the EGL. A recent study indicates that Numb also participates in coordinated endocytosis of the Shh receptors Boc and Ptch1, and thereby enables non-canonical Shh signaling (Ferent et al., 2019). Therefore, inactivation of aPKC via Eya1 might present a mechanism to promote symmetric divisions of GCPs, preserve canonical Shh signaling, and inhibit migration out of the mitogenic niche that is provided by the EGL.

It has been shown that the highly conserved C-terminal domain (CTD) of Eya family members contains Tyr phosphatase activity (Jemc and Rebay, 2007; Li et al., 2003; Rayapureddi et al., 2003; Tootle et al., 2003). While the less conserved N-terminal domain (NTD) does not contain a sequence motif that shares homology to any other known phosphatase, it has repeatedly been documented that the NTD of Eya members harbors Ser/Thr phosphatase activity (Li et al., 2017; Liu et al., 2012; Okabe et al., 2009; Xu et al., 2014). A recent study suggests that Ser/Thr phosphatase activity reported for Eya3 reflects a close association between the Eya NTD and the PP2A enzyme (Zhang et al., 2018), rather than direct activity of Eya3 itself. Here, we suggest that the CTD of both murine and human Eya1 is essential for full Thr phosphatase activity, as alterations of the D327 residue within the CTD domain reduce Eya1 activity on Thr-containing peptides, albeit to a lesser extent than its activity on Tyr-containing peptides (Li et al., 2017). In conclusion, our data indicates Eya1 promotes active Shh signaling-driven symmetric division of GCPs during early postnatal development (Fig. 7). Eya1 dephosphorylates aPKC, a component of the PAR complex. Consequent decreased phosphorylation of Numb prevents asymmetric distribution of NUMB in dividing GCPs. Therefore, both daughter cells continue into the next mitotic cell cycle. However, as development proceeds, decreased levels of Shh and Eya1 (See Figure 4B) result in asymmetric division of GCPs, and two distinct cell fates for daughters (Fig. 7). Given the Shh dysregulation in various cancers including medulloblastoma, these data suggest that Eya may be an important contributor to tumor initiation and/or rate of tumor growth.

**Fig. 7.**
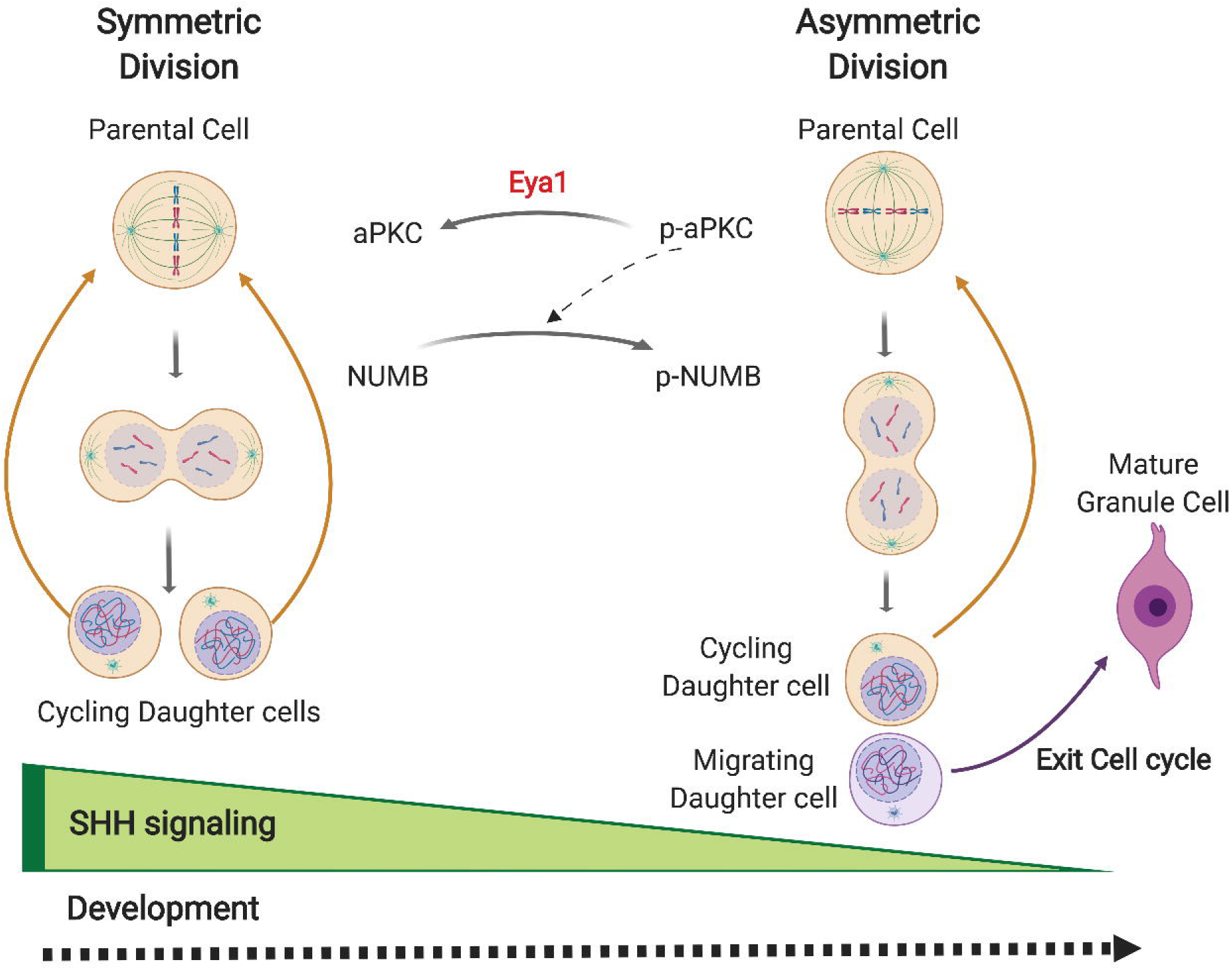
Eya1 promotes SHH-driven symmetric division in GCPs during cerebellar development. A model of Eya1 and SHH regulation of symmetric division for GCPs. During development, SHH signaling is highly active, thereby promoting symmetric division. Eya1 de-phosphorylates aPKC, which leads to reduced phosphorylation of Numb and equal distribution of Numb to the two daughter cells. Therefore, the daughter cells both remain in the EGL and re-enter the cell cycle. As development proceeds, Shh and Eya1 expression decrease, and therefore GCPs divide asymmetrically, producing one daughter cell that exits the cell cycle and migrates to the IGL, and one that reenters the mitotic cycle.

---

**Fig. S1. Spindle orientation of GCPs is developmentally regulated.** (A) Immunostainings for phospho-Histone H3 (pH3) of cerebellar sections from P3 and P6 wild type mice and representative images for vertically (left) and horizontally (right) dividing GCPs in the EGL. (B) Quantification of the distribution of angles of division of GCPs during the course of postnatal development. (C) Angles of division of GCPs in relation to the distance from the pial surface.

**Fig. S2. Active Shh signaling is recapitulated in MEF cells by SAG induction.** Mouse embryonic fibroblast (MEF) cells were induced with SAG (300nM) for 24h and *Gli1* mRNA levels were quantified using quantitative real time PCR. \*\**P*<0.01, unpaired *t* test. Data are mean±s.e.m.

## Materials and Methods

### Animal Studies

All experimental procedures were done in accordance with National Institutes of Health (NIH) guidelines and were approved by the Dana-Farber Cancer Institute’s Institutional Anima Care and Use Committee. *Eya1*^−/−^ mice were obtained from Pin-Xian Xu (Xu et al., 1999). *Shh^ala^* (Chan et al., 2009), *Ptch1*^+/−^ (Goodrich et al., 1997), loxP-Stop-loxP-EGFP (*GFP*^*Fl*/+^) (Mao et al., 2001), and *Atoh-1creER* (Machold and Fishell, 2005) mice were obtained from Jackson Laboratories.

### Real-time imaging in organotypic slices

Atoh1-Cre-Estrogen Receptor mice (Atoh1-creER) were crossed with *GFP^Fl/Fl^* (loxP-Stop-loxP-EGFP) mice to generate *Atoh1-creER:GFP*^*Fl*/+^ mice. Neonatal pups (the day when pups were born is designated as P1) were injected intraperitoneally with 10 μl 10 mg/ml tamoxifen. Thirty hours after tamoxifen injection (P3), the pups were dissected to isolate the cerebellum and other regions of the brain were used to check GFP expression by fluorescence microscopy. Cerebella were embedded in 2% low-melting agarose and cut into 200 μm slice. Cerebellar slices were cultured on tissue culture inserts (4 slice per insert) in 6-well plates with phenol-free GCP growth medium at 37°C for 2 hours for recovery before imaging. A Nikon Perfect Focus microscope (Ti Eclipse; Nikon) equipped with a CO2 controlled chamber (5%CO2 and 37°C), a CCD video camera and a far distance 20x objective were used to capture the images on multiple positions with 5 μm-thickness z stacks (1 μm intervals between z-stack) every 45 minutes for the next 72 hours. Time-lapse experiments were analyzed using NLS-Elements software (Nikon). We then analyzed the videos, to identify individual cells that divide within the first 24 hours. We analyzed whether both of the GFP labeled daughter cells remain in the EGL where they will continue to divide (vertical division), or whether one divides and the other migrates to the IGL (horizontal division), or whether both daughters migrate to the IGL.

### Immunohistochemistry

For cryostat sections, mouse brains or cerebella were fixed in 4% paraformaldehyde overnight at 4°C followed by cryoprotection in 30% sucrose until sinking to the bottom. Sagittal and coronal sections (10 μm) were prepared using a cryostat (Leica Micro-systems). For paraffin sections, mouse brains were fixed in 4% paraformaldehyde overnight, embedded in paraffin using standard protocols, and then cut (5 μm sections). For cerebellar organotypic slices, slices were prepared with a Leica vibratome followed by fixation with ice-cold methanol at −20°C for overnight. Slices were washed 3 times and rehydrated in PBS overnight. All brain sections were blocked and permeabilized with blocking/permeabilization solution (5% normal goat serum, 0.1% Triton X-100, and 5% BSA in PBS) for 1 hour at room temperature or at 4°C overnight, followed by incubation with primary antibody at 4°C overnight. After incubation with primary antibodies, sections were washed in PBS, followed by the incubation with appropriate Alexa Fluor 546, 647, or 488-labeled secondary antibodies (Invitrogen, 1:400) and DAPI counterstain. Images were acquired on either a Nikon upright fluorescence microscope and NLS-Element software (Nikon) or confocal microscope (SP5; Leica) using 20X and 40X (Plan Neofluar NA 1.3 oil immersion) objectives and LAS software (Leica). Fiji software was used for image processing (Gaussian blur) and data analysis (spindle orientation measurement).

### Quantitative analyses of asymmetric protein distribution

Cryosections from postnatal day 3, wild type cerebella were stained with Vybrant^®^ CM-DiI (Invitrogen) for 20 minutes at room temperature to mark the cell boundary, then post-fixed with 4% PFA. After washing with PBS, CM-DiI-labeling sections were blocked and permeabilized with blocking solution (5% NGS, 1% BSA, 0.1% Triton X-100 in PBS) for 1 hour and followed by co-immunostaining with antibodies to phospho-histone H3, to recognize mitotic cells and visualize mitotic orientation and with antibodies to symmetry marker LGN (GSPM2) or Numb, followed by DAPI counterstaining. Images were captured by confocal microscopy (Leica SP5). We examined all phospho-histone H3 positive cells in anaphase or telophase and accurately defined the boundaries of individual GCPs in the crowded EGL using CM-DiI. We further used phospho-histone H3 and DAPI to define the mitotic plane. We then integrated the signal for LGN or Numb across the Z-axis of the two incipient daughter cells. The “Asymmetry Index” is the absolute value of signal in Daughter A minus signal value in Daughter B/Signal in Daughter A plus signal in Daughter B. Complete symmetry gives an index of 0, while complete asymmetry gives an index of 1. We carried out this analysis both for integrated and mean measurements of our markers. A second individual determined whether the mitosis is perpendicular or parallel in nature based on the orientation of the chromosomes separating in anaphase or telophase (visualized with phospho-Histone H3 and with DAPI stain of DNA). Orientation was readily apparent in most anaphase/telophase cells. We also assessed a third category, of indeterminate/intermediate orientation, which encompasses cells that are in early stages of mitosis and those with oblique orientation. As a control for the distribution specificity of particular symmetry marker LGN and Numb in dividing cells, we also analyzed the Asymmetry Index for GAPDH.

### Measurement of mitotic spindle orientation

For staining using phospho-Histone H3, a line was drawn perpendicular to the mitotic spindle to best represent the plane of division. A second line perpendicular to the pial surface was drawn, and the angle between those lines was measured. For all stainings using acetylated α-tubulin and γ-Tubulin, a line was drawn between two centrosomes to mark the mitotic spindle. A second line between two points of the pial surface on the both ends of the fissure was drawn to act as a reference line for scoring the spindle orientation. For organotypic slices, we defined x, y, and z coordinates of the two centrosomes and five points on the pial surface of the 3D-rendered cerebellar slices. These five points were used to determine the best-fitting plane by orthogonal distance regression. Then, we calculated the angle between the line connecting two centrosomes and line for the best-fitting plane. The angle α of mitotic spindle orientation is calculated as 90° plus the angle. To estimate the uncertainty of each spindle orientation angle, we repeated this calculation for all possible combinations using only four out of the five points within the plane and determine the angular SD of the resulting normal lines. All angles were scaled to correspond to a range between 0° (vertical cell division) and 90° (horizontal cell division). Three pairs of control littler mates and mutants were analyzed. Comparable locations (midpoint of primary, secondary and tertiary fissures within vermis) were assessed.

### Cell culture and constructs

Mouse embryonic fibroblast (MEF) cultures from *Eya1*^−/−^ and wild type littermates were generated and cultured as previously described (Jozefczuk et al., 2012). For Shh activation, MEF cells were cultured for 24 hours in MEF media with reduced serum (0.5%) in the presence of 300 nM SAG. HEK293T cells were obtained from American Type Culture Collection and cultured according to their recommendations.

### Peptides and plasmids

All peptides were custom synthesized by the Tufts University Core Facility for Peptide Synthesis. All peptides were purified by high-performance liquid chromatography, and all purified products were analyzed by mass spectrometry. The purity of each peptide was greater than 95%. The coding sequence of full-length Eya1 was cloned into pcDNA3.1(+)-KozakHAHA as previously described (Eisner et al., 2015) for transfection into HEK293T cells. For protein expression in S2 cells, full-length Eya1 was cloned into a custom-modified version of the pFastBac vector HT A (Thermo Fisher) carrying the sequences for His and GST tags. For the construction of the D327A Eya1 mutant, we generated a single nucleotide exchange using the QuikChange II Site-Directed Mutagenesis Kit (Agilent).

### Protein expression and purification

Full-length Eya1 wild type and mutant D328A protein in a GST-fusion format were generated using the Bac-to-Bac Baculovirus Expression system (ThermoFisher Scientific) and expressed in Sf9 insect cells. Cells were harvested 65-72 hours postinfection and resuspended with lysis buffer (1X PBS, 10% Glycerol, 1 mM TCEP).

Resuspended cell pellets were disrupted by sonication. After sonication, lysed cells were spun down at 40000 rpm for 2 hours. Supernatants were applied to GST sepharose beads (GE Lifescience). After washing with PBS, bound proteins were eluted with GST Elution Buffer (50 mM Tris-HCl pH8.0, 150 mM NaCl, 10 mM Glutathione, 2 mM TCEP, 10 mM MgCl_2_). To remove GST tag, eluents were incubated with Tabacco etch virus protease for 6 hours on ice. After cleavage, eluents were applied to a HiTrap Q column (GE Lifescience) and washed with Buffer A (Buffer A. 50 mM Tris-HCl pH8.0, 2 mM TCEP, 10 mM MgCl_2_). Eya1 protein was eluted with a 0 – 25% gradient of Buffer B (Buffer B. 50 mM Tris-HCl pH8.0, 2 M NaCl, 2 mM TCEP, 10 mM MgCl_2_). Eya1 protein was concentrated to approximately 2 mg/ml using an Amicon Ultra concentrator (30 MWCO, Millipore) and further purified by size-exclusion chromatography on a Superdex 200 Increase 10/300 column (GE Lifescience) in Buffer C (Buffer C. 40 mM HEPES pH 7.5, 150 mM NaCl, 10 mM MgCl_2_, 2 mM TCEP). Purified Eya1 protein was concentrated to 1mg/ml and frozen by liquid nitrogen for long-term storage.

### *In vitro* phosphatase assay

For peptide assays in half-area 96 well plates, 400 ng (unless specified otherwise) of purified Eya1 protein was incubated in a final volume of 20 μl with 1 mM peptide in a buffer containing 60 mM HEPES (pH7.5), 75 mM NaCl, 75 mM KCl, 5 mM MgCl_2_, 1 mM EDTA, and 1 mM DTT at 37°C for 2 hours. The released phosphate was quantified using the Malachite Green Phosphate Assay Kit (Sigma-Aldrich). Duplicates values were background corrected and compared to the internal phosphate standard.

### Quantitative real-time RT-PCR

RNA was isolated using the RNeasy Mini Kit (Qiagen). Reverse transcription was performed using the High Capacity cDNA Reverse Transcription Kit (Applied Biosystems) according to manufacturer’s specifications. qRT-PCR was performed using Taqman gene expression assays with the following Taqman probes: *Eya1* Mm01239746_m1, *Shh* Mm00436527_m1, *Ptch1* Mm00970977_m1, *Smo* Mm01162710_m1, *Sufu* Mm00489385_m1, and *Gli1* Mm00494645_m1. Values were normalied to *Gapdh* levels.

### Co-immunoprecipitations and western blotting

For all assays, protein was extracted using RIPA buffer. For co-immunoprecipitations, we used Dynabeads Protein G (Invitrogen) according to manufacturer’s protocol and optimized antibody concentrations. For western blots, protein lysates were separated electrophoretically in NuPAGE 4 – 20 % Bis-Tris gels (Thermo Fisher) and blotted on a PVDF membrane (Millipore) using standard procedures. Optimal concentrations for all antibodies were evaluated empirically. Protein bands were visualized using horse radish peroxidase (HRP)-coupled secondary antibodies (BioRad) and SuperSignal West Dura solution as substrate (Thermofisher). Image acquisition was done HyBlot CL autoradiography film (Denville Scientific).

### Statistical Analysis

Significance threshold for *P* value was 0.05 (\**P* < 0.05, \*\**P* < 0.01, \*\*\**P* < 0.001, \*\*\*\**P* < 0.0001). For comparison of two groups, we used a Student t test. We used a two-way ANOVA with Bonferroni correction for multiple comparisons to compare data from several groups. For comparison of fractions, Fisher’s exact test was used.

## Author Contributions

Conceptualization: R.A.S, K.J.N, J.A.C, P.C.Z, D.J.M; Methodology and formal analysis: D.J.M, P.C.Z, S.M.C, M.P.-M, K.J.R, K.J.N, R.A.S; Investigation: D.J.M, P.C.Z, S.M.C. M.P.-M, K.J.N, J.A, X.Z, E.P. K.J.N, R.A.S; Resources: P.-X.X, E.P, M.J.E, J.A.C; Writing – original draft: P.C.Z, D.J.M, R.A.S; Writing – review & editing: D.J.M, M.P.-M. R.A.S, and All authors; Visualization: D.J.M, P.C.Z, G.H; Funding acquisition: R.A.S, K.J.N, J.A.C, D.J.M

## Acknowledgements

We thank CD Stiles and all the members of the Segal lab for helpful suggestions.

